# PD-1 signaling is essential for the early accumulation of HBV-specific CD8+ T cells during HBV infection

**DOI:** 10.1101/2025.09.17.676789

**Authors:** Xiaoqing Zeng, Wen Pan, Ziwei Li, Zhaoli Liu, Hongming Huang, Xuecheng Yang, Kathrin Sutter, Mengji Lu, Ulf Dittmer, Gennadiy Zelinskyy, Xin Zheng, Dongliang Yang, Patrick T.F. Kennedy, Yanqin Du, Jia Liu

## Abstract

**Background & Aims:** Programmed cell death Protein 1(PD-1) is one of the key inhibitory receptors that regulates CD8 T cell exhaustion during chronic viral infection. However, the role of PD-1 in modulating effector T cell differentiation and function during early phase of hepatitis B virus (HBV) infection remains poorly defined.

**Methods:** Using adoptive transfer of wild-type or PD-1 KO HBV-specific TCR transgenic (C93-TCRtg) CD8⁺ T cells, flow cytometry, and transcriptome sequencing, we systematically profiled early PD-1 dynamics in splenic and intrahepatic T cells and evaluated how PD-1 deficiency shapes anti-HBV T cell responses in acute self-resolving (AR) and chronic (CH) HBV mouse models.

**Results:** Acute HBV infection induced sustained PD-1 upregulation on intrahepatic/splenic CD8⁺ T cells, which predominantly exhibited an effector phenotype. PD-1⁺ cells displayed enhanced proliferation, cytotoxicity, and effector cytokine production compared to PD-1⁻ counterparts. Adoptive transfer of PD-1 KO C93-TCRtg CD8⁺ T cells resulted in significantly impaired early expansion in both spleen and liver during acute infection, accompanied by reduced granzyme B expression while maintaining IFN-γ, TNF-α, or IL-2 production. Transcriptomic analysis revealed that PD-1 deficiency was associated with the upregulation of apoptosis gene signatures. In chronic HBV infection, PD-1 KO cells failed to expand long-term and exhibited exacerbated exhaustion evidenced by a loss of IFN-γ production.

**Conclusions:** PD-1 is essential for the early expansion and survival of HBV-specific CD8⁺ T cells during acute infection and does not suppress their effector cytokine function. Its absence impairs proliferation and promotes exhaustion of HBV-specific CD8⁺ T cell in chronic infections.

**Impact and implications:** Our study redefines PD-1’s role in acute HBV infection. Rather than merely constraining effector function, PD-1 is essential for the early expansion and survival of HBV-specific CD8^+^ T cells, acting as a safeguard against activation-induced apoptosis. The context-dependent functions position PD-1 as a critical rheostat calibrating the magnitude and quality of the antiviral CD8^+^ T cell response, with significant implications for immunotherapeutic strategies targeting this pathway in hepatitis B.

## Introduction

Hepatitis B virus (HBV) infection affects billions of individuals worldwide, with a significant proportion of chronic HBV infection progressing to liver cirrhosis and hepatocellular carcinoma. Achieving cure for chronic HBV remains relatively challenging. The ultimate control and clearance of the virus depend on the host’s immune response against HBV, particularly the virus-specific CD8^+^ T cell response, which plays a crucial role in this process [1] . Studies have found that in patients with acute self-limited infections, an obvious proliferation of HBV-specific CD8^+^ effector T cells secreting interferon-gamma (IFN-γ) can be detected, while the HBV-specific CD8^+^ T cell response was often weakened or in a state of functional loss/exhaustion in patients with chronic hepatitis B (CHB) [2, 3].

A hallmark feature of these exhausted T cells is over-expression of immune checkpoint molecules, such as cytotoxic T-lymphocyte associated protein 4 (CTLA4) and programmed cell death protein 1 (PD-1), and programmed cell death protein ligand1(PD-L1) [4]. Numerous ex vivo studies utilizing peripheral blood mononuclear cells (PBMCs) collected from patients with CHB have demonstrated that PD-1/PD-L1 blockade can enhance the HBV-specific CD8^+^ T cell response [5–7]. In 2014, our group for the first time reported the effects of in vivo administration of PD-L1 blocking antibodies on enhancing virus-specific CD8^+^ T cell immunity in woodchucks infected with chronic woodchuck hepatitis virus (WHV), a classic animal model for HBV infection research [8]. Our findings demonstrated that anti-PD-L1 blockade monotherapy was insufficient to rescue WHV-specific T cell function; however, when combined with antiviral treatment and therapeutic vaccination, it significantly enhanced WHV-specific CD8^+^ T cell immunity. The triple-therapy strategy led to sustained immunological control of viral infection after antivirals withdrawal, woodchuck hepatitis surface antigen (WHsAg) seroconversion and even complete viral clearance in some treated animals [8].

Although high expression of PD-1 is regarded as a typical surface marker of exhausted T cells, its upregulation actually begins following the initial recognition of specific antigens by T cell receptor (TCR) [9]. Therefore, PD-1 is also expressed during the early phases of T cell activation when naive CD8^+^ T cells differentiate into effector cells. In the acute phase of viral infections, such as lymphocytic choriomeningitis virus (LCMV), Friend virus (FV), hepatitis C virus (HCV), and HBV, upregulation of PD-1 on virus-specific T cells can be detected [10–15]. Study utilizing the FV model has shown that CD8+ T cells expressing PD-1 are activated effector T cells, not exhausted T cells during acute infection [15]. During acute LCMV or FV infection, blocking the binding of PD-1 and its ligand PD-L1 with antibodies can further enhance the activation and effector function of effector T cells [11, 15]. Similarly, in the acute phase of HBV infection, upregulation of PD-1 expression on peripheral blood CD8^+^ T cells has also been observed[13], but the dynamic characteristics of PD-1 expression on T cells in the liver and peripheral immune organs are currently unclear. While PD-1 is a well-established suppressor of T cell function in chronic HBV, its paradoxical sustained upregulation on functional effector CD8⁺ T cells during acute self-resolving infection suggests a context-dependent role that remains mechanistically unresolved. Understanding whether PD-1 acts as an activator or inhibitor in acute HBV is critical for designing checkpoint-based immunotherapies, as premature blockade may inadvertently impair viral clearance.

In this study, we employed HBV replication mouse models to longitudinally characterize PD-1 expression patterns on CD8^+^ T cells in the spleen and liver across different viral clearance outcomes following HBV exposure. Furthermore, through adoptive transfer of PD-1-deficient HBV-specific CD8^+^ T cells, we investigated the early impact of PD-1 deficiency on the development of HBV-specific CD8^+^ T cell responses during HBV infection. Using these approaches, we found that PD-1 is required for the early expansion of HBV-specific CD8^+^ T cells and does not suppress their effector functions during the acute phase of HBV infection.

## Materials and methods

### Mice

Male wild-type CD45.2 C57BL/6 mice (6 to 8 weeks old) were purchased from Hunan Slack King Laboratory Animal Co., Ltd. PD-1^-/-^ mice were kindly provided by Prof. Ran He from Tongji Hospital, Tongji Medical College, Huazhong University of Science and Technology. CD45.1 congenic mice were provided by Professor Ulf Dittmer from University of Duisburg-Essen. Core93 TCR-transgene (C93-TCR-tg) mice were purchased from the Jackson laboratory. PD-1^-/-^ C93-TCRtg mice were generated by crossing PD-1 knockout mice with C93-TCRtg mice to obtain HBcAg-specific CD8+ T cells lacking PD-1 expression. All mice were maintained under specific pathogen-free (SPF) conditions in the Animal Care Center of Tongji Medical College according to the Guidelines of the National Institute of Health for Animal Care and Use. The protocols and procedures employed were ethically reviewed and approved by the institutional Animal Care and Use Committee at Tongji Medical College, Huazhong University of Science and Technology.

### HBV replicating mouse model

HBV replication plasmids pSM2 used in this study were provided by Dr. Hans Will, Heinrich-Pette-Institute, Hamburg, Germany. An acute HBV replication mouse model was established by hydrodynamic injection (HDI) of 10 μg pSM2 plasmid as described before [16]. Briefly, HBV plasmid was injected into tail vein of male 6–8 weeks old mice in a volume of normal saline (NS) equivalent to 0.1 mL/g of the mouse body weight within 5 s. Chronic HBV replication mouse model was established by HDI of 6μg pAAV/HBV1.2 plasmids or intravenous injection (i.v) of recombinant adeno-associated virus type 8 carrying the 1.3-mer wild-type HBV genome (rAAV8-1.3HBV) (FivePlus Molecular Medicine Institute, Beijing). A total of 5 × 10^10^ viral genomes/200 μl virus were injected into the tail vein of each C57BL/6 mouse.

### Detection of serological HBV infection markers

Serum hepatitis B surface antigen (HBsAg), hepatitis B surface antibody (HBsAb), and hepatitis B e antigen (HBeAg) were detected using enzyme-linked immunosorbent assays (ELISAs) (KHB, Shanghai, China) following the manufacturer’s instructions, after tenfold dilution with phosphate-buffered saline (PBS). Serum HBV DNA levels were detected by real-time PCR using commercial reagents (Sansure, Hunan, China), according to the manufacturer’s instructions.

### Cell isolation

Single-cell suspensions of murine splenocytes were obtained by ground on a 70-μm filter. The mouse liver-infiltrating lymphocytes (LILs) were isolated through Percoll density centrifugation after the livers were perfused with PBS via the portal vein as described previously [17]. In brief, mouse livers were perfused with 10 ml PBS immediately after sacrifice. The livers were then homogenized, and cell pellet was collected. Then cell pellet was resuspended in 40% Percoll and centrifuged at 1000 g without break. After removing the debris and hepatocytes on the top layer, LILs in the pellet were collected, washed, and subjected to further analysis. In addition, preparation of single-cell suspensions of splenocytes was performed by ground on a 70-µm filter. Pellet was then incubated with 5 ml red blood cell lysis buffer for 5 min. After washing twice, splenocytes in the pellet were collected and subjected to further analysis.

### Cell sorting and adoptive transfer

The splenic CD8 +T cells were isolated by the CD8+T Cell Isolation Kit (130-104-075, Milteyni Biotec) according to the manufacturer’s instructions. The purity of isolated CD8 + T cells and B cells was > 95% after isolation. Different number of purified CD8+T cells were transferred into recipient mice by i.v at indicated time points.

### Detection of HBV-specific T cell response

Freshly isolated LILs or splenocytes were first incubated with anti-CD16/32 rat anti-mouse antibody (eBioscience), and then stained with HBV core Tetramer-MGLKFRQL-APC (Institute of Medical Biology, Co., Ltd.) for 60 min at 4°C. After washing, the cells were stained with anti-CD8 and fixable viability dye (FVD) (eBioscience) and detected on flow cytometry.

For detecting the function of HBV-specific T cells, freshly isolated LILs or splenocytes were stimulated with 10 μg/ml H-2K^b^-restricted HBcAg-derived CD8^+^ T cell epitope peptide core93 to 100 (MGLKFRQL) at 37°C for 5 h in the presence of 1 μg/ml anti-CD28 antibody (eBioscience), 1 μg/ml brefeldin A, and 1 μg/ml monensin (eBioscience). The cells were primarily stained with anti-CD8 and fixable viability dye eFluor 506 (FVD) (eBioscience), followed by intracellular cytokine staining (ICS) with monoclonal antibodies against interleukin-2 [IL-2], interferon-gamma [IFN-γ], and tumor necrosis factor-alpha [TNF-α] (all from Biolegend).

### Flow cytometry

Surface staining was performed by incubating the cells for 30 min in the dark at 4 °C with the desired antibody combination. For intracellular staining, the cells were fixed and permeabilized using the intracellular fixation and permeabilization buffer set (Biolegend) for 30 min and then stained with the desired antibodies. For staining of Ki-67, Eomes, T-bet and Tox, cells were fixed and permeabilized using the fixation/permeabilization buffer set (eBioscience) for 45 min at room temperature and then stained with the desired antibody combinations. The monoclonal antibodies used are listed in the **Table S1**. The data were acquired on FACS Canto II Flow cytometer (BD Biosciences) and analyzed with FlowJo 10 software (TreeStar, Ashland, USA). The cell debris and dead cells were excluded from the analysis based on forward/ sideward scatter signals and fixable viability dye eFluor 506 (FVD) (eBioscience) staining.

### In Vitro stimulation of murine lymphocytes

Freshly isolated splenocytes from Wild type (WT) or PD-1 knockout (KO) HBcAg-specific TCR transgenic (C93-TCRtg) mice were stained with CellTrace™ CSFE according to the manufacturer’s instruction (Thermo Fisher Scientific), and the cells were stimulated with 10 μg/ml HBV peptide core93-100 (MGLKFRQL) in the presence of 1 μg/ml anti-CD28 (eBioscience) at 37 °C in 5% CO_2_ for 96 h. The secretion of IFN-γ, TNF-α in the supernatant were measured by CBA methods according to the manufacturer’s instruction (Bioscience, BD).

### Bioinformatic analysis

High throughput sequencing and data analysis were performed by BGI Huada. Differentially expressed genes were selected using the following criteria: fold change >2 (log2 ratio ≥ ±1), false discovery rate (FDR) <0.001. Gene set enrichment analysis (GSEA) was performed by GSEA 4.4.0 software. All gene sets from Hallmarks (40 gene sets), C2 (6290 gene sets) and C7 (5219 gene sets) databases from MsigDB were tested for a total of 11559 gene sets.

### Statistical analysis

Statistical analyses were performed using GraphPad Prism 8 software. The differences between two groups were evaluated using *t* test. The statistical differences among multiple groups were analyzed by one-way ANOVA followed by Turkey’s multiple comparisons test. The statistical differences of viral indicators were analyzed by repeated measures analysis. Data are presented as the mean ± standard error (SEM) and differences were considered statistically significant at *p* < 0.05. Significance is denoted with asterisks (\**p*< 0.05, \*\**p*< 0.01, \*\*\**p*< 0.001).

## Results

### Prominent and sustained PD-1 upregulation on CD8^+^ T cells in the spleen and liver characterizes acute self-resolving HBV replication

To characterize the role of PD-1^+^CD8^+^ T cells during the acute phase of HBV infection, we used the well-established acute and chronic HBV replication mouse models by HDI of HBV replication plasmids pSM2 and pAAV/HBV1.2 into male C57BL/6 mice. HBV markers, including HBsAg and HBV DNA, were monitored for 28 days post-injection (dpi) (**Fig. 1A**). Consistent with our previous study [17], all mice with acute self-resolving HBV replication (AR) were negative for serum HBsAg and HBV DNA at 21 dpi. In contrast, over 80% of the mice with chronic HBV replication (CH) remained positive for serum HBsAg and HBV DNA at 28 dpi (**Fig. 1B**). We then conducted a longitudinal analysis of PD-1 expression on CD8^+^ T cells in the liver and spleen following HBV exposure. Significantly increased frequencies of PD-1 expressing CD8+ T cells in the liver were observed at 14, 21, and 28 dpi in AR mice, whereas no such increase was observed in CH mice. CD8^+^ T cells in the spleen of AR mice also demonstrated sustained increase in PD-1 expression throughout the same period (14–28 dpi). In contrast, CH mice showed only a transient PD-1 upregulation on CD8^+^ T cells in the spleen at 14 dpi, with levels returning to baseline thereafter (**Fig. 1C**). Next, we investigated PD-1 expression dynamics in HBV core-specific CD8⁺ T cells. Core93-specific CD8^+^ T cells in the liver of AR mice exhibited a progressive increase in PD-1+ cells, peaking at 91.1% by 21 dpi. Moreover, PD-1^+^ cell proportions in core93-specific CD8+ T cells consistently surpassed those in non-core93-specific CD8⁺ T cells across all time points analyzed (**Fig. 1D**). Taken together, these results demonstrate distinct temporal and spatial patterns of PD-1 expression on CD8^+^ T cells during the acute phase of HBV infection between AR and CH models, with AR marked by strong and sustained intrahepatic/splenic PD-1 upregulation on CD8^+^ T cells, contrasting sharply with weak and transient PD-1 induction in chronic infection.

**Figure 1.**
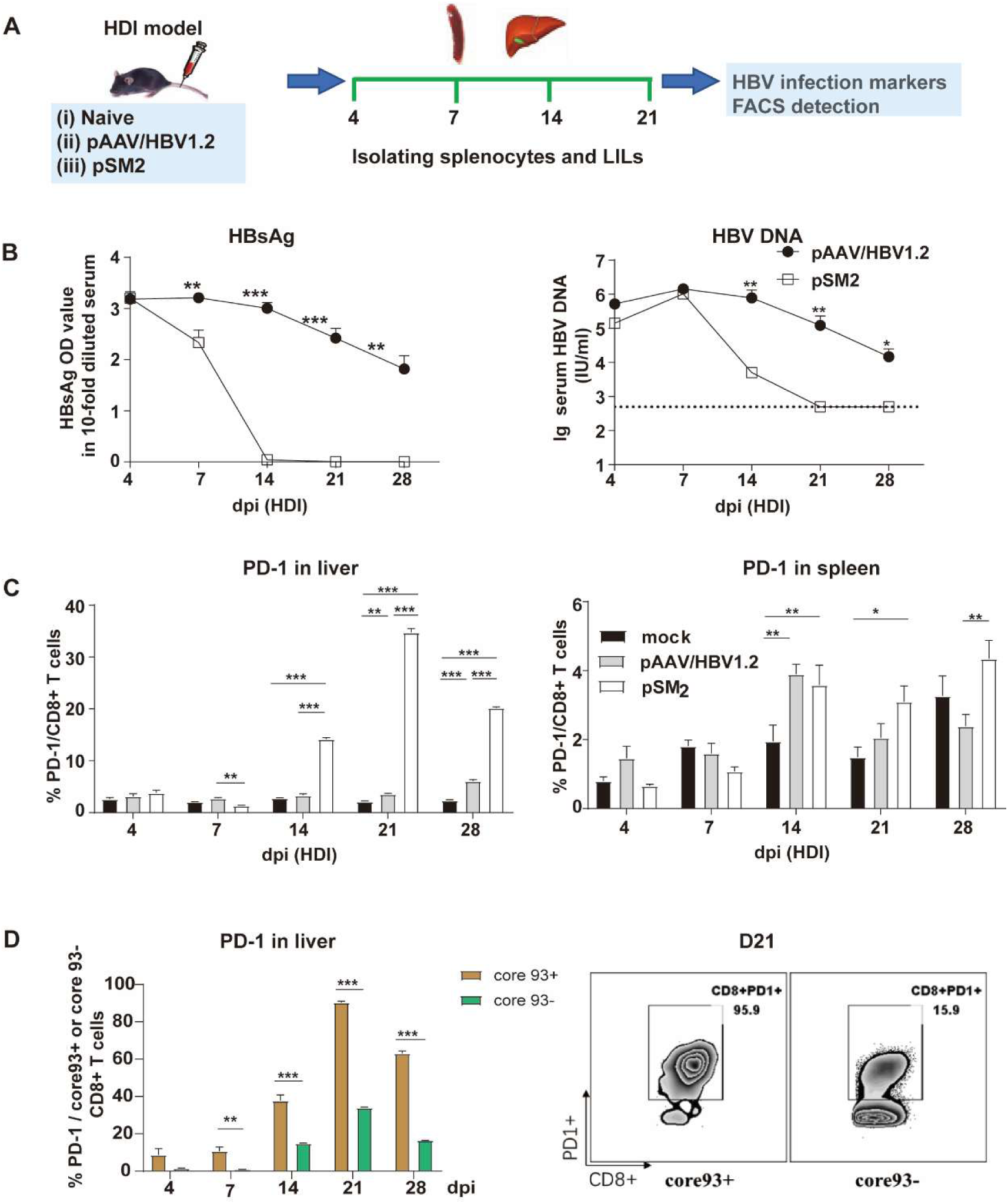
Expansion of PD-1^+^ CD8^+^ T cells in the liver and spleen in the acute self-resolving HBV replication mouse model. (A) Schematic overview of the experiment setup of AR and CH model: mice were injected with pSM2 plasmid, pAAV/HBV1.2 plasmid or mock control by HDI. LILs and splenocyte were isolated at indicated time points. (B) The kinetics of serum HBsAg and HBV DNA were determined. (C) The expression of PD-1 on CD8+ T cells in liver and spleen was illustrated. (D) The representative staining and the expression of PD-1 in HBc-specific (core 93+) CD8+ T cells and non-HBc-specifc (core 93-) CD8+ T cells were shown. Data are depicted as arithmetic means ±SEM. The statistical differences among multiple groups were analyzed by one-way ANOVA followed by Turkey’s multiple comparisons test. The statistical differences of viral indicators were analyzed by repeated measures analysis. Five to six mice were analyzed per group, and at least two independent experiments were performed. \**p*<0.05, \*\**p*< 0.01, \*\*\**p*<0.001. *AR*, acute self-resolving HBV replication; *CH,* chronic HBV replication; *dpi,* days postinfection; *HDI,* hydrodynamic injection; *LIL,* liver infiltrating lymphocytes.

### PD-1^+^CD8^+^ T cells are dominated by an effector phenotype during the acute phase **of self-resolving HBV replication.**

Given the observed significant upregulation of PD-1 expression on CD8^+^ T cells in AR mice, we next performed a comparative analysis of the differentiation, proliferation, and effector function of PD-1^+^ CD8^+^ T cells. The differentiation status of T cells was assessed based on the expression of CD44 and CD62L, as previously described [18]. Effector and effector memory T cell (T_E_/T_EM,_ CD44^+^CD62L^-^) phenotype predominated in PD-1^+^CD8^+^T cell population, showing significantly higher frequencies compared to PD-1^-^CD8^+^T cells at 5 all investigated time points both in the spleen and liver (**Fig 2A)**. Notably, nearly all PD-1+ cells had differentiated into an effector T cell phenotype from the outset in the liver (**Fig 2A)**. PD-1^+^ cells also exhibited a highly active proliferative state, as evidenced by sustained high expression of the proliferation marker Ki-67 compared to PD-1^-^ cells in the spleen (**Fig 2B)**. Next, we elucidated the antiviral function of PD-1^+^ T cells by measuring the expression of granzyme B and effector cytokines in both PD-1^+^ and PD-1^-^CD8^+^ T cells. Significant upregulation of granzyme B were observed in PD-1+CD8+T cells compared to PD-1^-^CD8^+^ T cells in both the liver and spleen (**Fig. 2C)**. In the spleen, PD-1^+^CD8^+^ T cells exhibited significantly stronger cytokine secretion capacity (IFN-γ, IL-2, and TNF-α) compared to PD-1^-^CD8^+^ T cells upon stimulation at all time points tested (4, 7, 14, and 28 dpi) (**Fig. 2D**). In contrast, liver-specific differences were predominantly confined to the early stages of infection. During 4–14 dpi in AR mice, PD-1^+^CD8^+^ T cells in the liver produced markedly higher levels of IFN-γ, IL-2, and TNF-α than their PD-1-counterparts. However, this functional disparity diminished by 21 dpi, with no significant differences in IFN-γ or IL-2 production observed between PD-1+ and PD-1-subsets at 21 and 28 dpi, although TNF-α production remained elevated (**Fig. 2D**). Collectively, these results suggest that PD-1^+^CD8^+^ T cells predominantly exhibit an effector T cell phenotype and maintain sustained proliferative activity and antiviral effector functional capacity in acute phase of self-resolving HBV replication.

**Figure 2.**
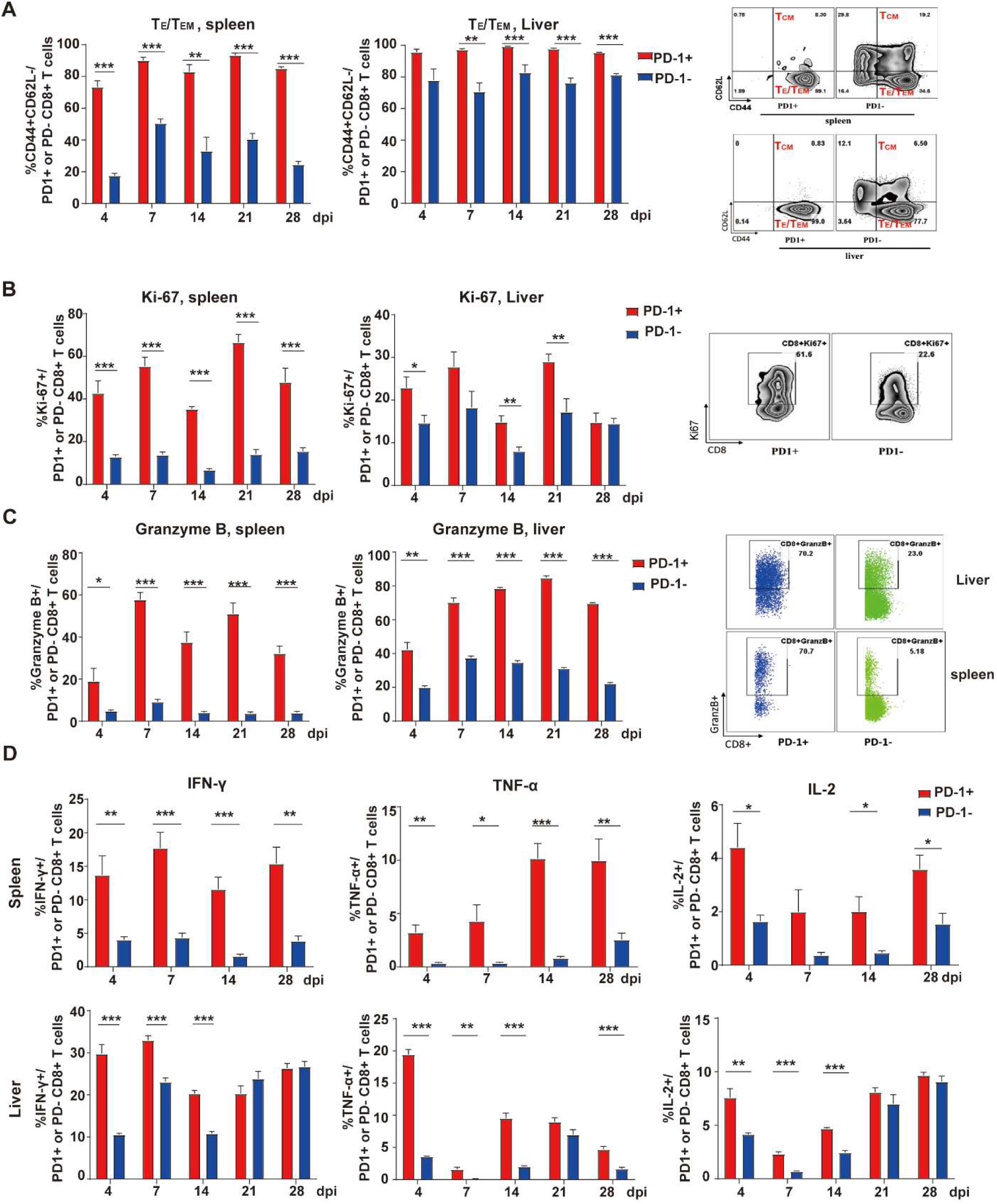
PD-1^+^ CD8^+^ T cells are dominated by T_E_/T_EM_ phenotype and displayed antiviral function during the acute HBV replication model. Mice were injected with pSM2 plasmid by HDI. LILs and splenocytes were isolated at indicated time points. (A) The representative staining (right panel) and the summary (left panel) of the differentiation on PD-1^+^ or PD-1^-^ CD8+ T cells were shown. (B) The expression of proliferation marker Ki-67 and (C) degranulation marker Granzyme B on PD-1+ or PD-1-CD8+ T cells were determined. (D) Cells were stimulated with CD3 and CD28 in the presence of Brefeldin A for 5h. The frequencies of IFN-γ, IL-2 and TNF-α out of PD-1^+^ or PD-1^-^ CD8+ T cells were displayed. Five mice were analyzed per group. Data are depicted as arithmetic means ± SEM. Differences between mouse groups were analyzed using unpaired Student’s *t* tests. \**p*< 0.05, \*\**p*< 0.01, \*\*\**p*< 0.001. *HDI,* hydrodynamic injection; *IFN-γ*, interferon-gamma; *IL-2*, interleukin-2; *TNF-α*, tumor necrosis factor-alpha.

### Transcriptomic profiling reveals attenuated apoptosis in PD-1^+^CD8^+^ T cells of AR **mice**

To further investigate the phenotypic characteristics of PD-1+CD8+ T cells, we isolated PD-1^+^CD8^+^ T cells and PD-1^-^CD8^+^ T cells from the livers of AR mice at 21dpi for transcriptome sequencing analysis (**Fig.3A**). A total of 3009 differentially expressed genes were identified between the two groups, with 1339 genes upregulated (red) and 1670 genes (blue) downregulated in PD-1^+^CD8^+^ T cells compared to PD-1^-^CD8^+^ T cells (**Fig.3B-C**). Notably, 140 of these differentially expressed genes were associated with cell growth and death pathway (**Fig.3D**). Among the top 20 differentially expressed genes, several additional inhibition markers, including CTLA-4, TIGIT and LAG3, were also significantly upregulated in PD-1^+^CD8^+^ T cells relative to PD-1^-^CD8^+^ T cells (**Fig.3E**). Protein-protein interaction analysis revealed that genes related to the cell cycle, cell growth and death, and immune function clustered together (**Fig.3F**). Critically, GSEA analysis displayed significant enrichment of apoptosis gene signature in PD1-CD8+ T cells compared to PD-1^+^ CD8^+^ T cells at 21dpi (**Fig.3G**). Collectively, these findings indicate that PD-1^+^ CD8^+^ T cells exhibit upregulation of other inhibition markers, and display less apoptosis gene enrichment.

**Figure 3.**
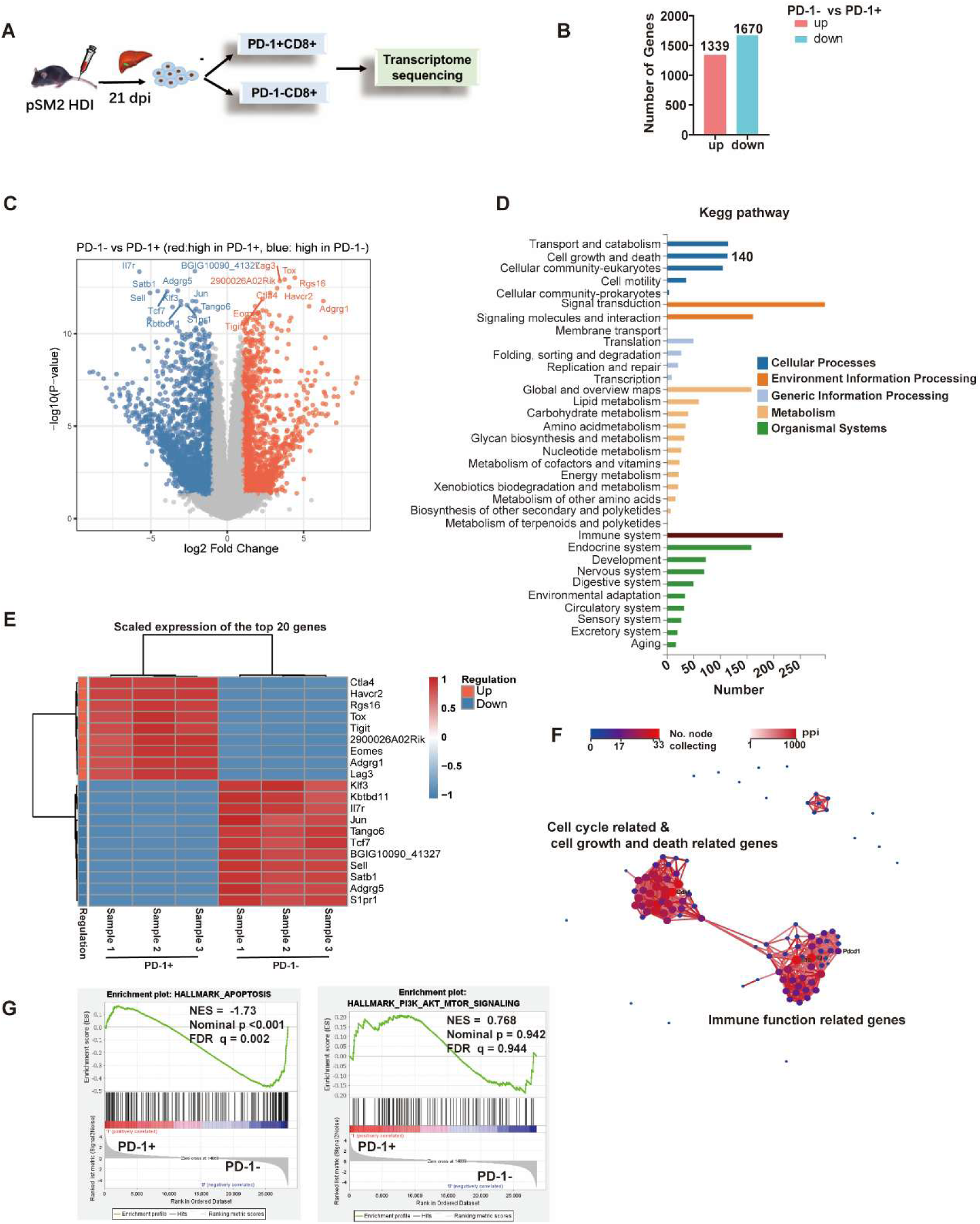
Differentially expressed genes between PD-1+ CD8+ T cells and PD-1-CD8+ T cells. (A) Schematic overview of the experiment setup: PD-1+ and PD-1-CD8+ T cells were isolated from liver at 21 dpi after HDI of pSM2 plasmid and then subsequently sent for transcription sequencing. (B) The number of differentially expressed genes between two groups was shown. (C) Volcano plots showed differentially expressed genes between two groups. The x axis shows the log2 Fold change (logFC) comparing PD-1+ with PD-1-CD8+ T cells, and the y axis shows the -log10 p value in the association. (D) KEGG pathway of different function was shown. (E) The top 20 differentially expressed genes between two groups were shown by heatmap (FDR<0.05). (F) Protein-protein interaction network analysis of cell cycle related, cell growth and death related genes as well as immune function related genes were shown. (G) GSEA of apoptosis genes and PI3K/Akt/mTOR signaling genes signatures of PD-1+ and PD-1-CD8+ T cells. There samples were used in each group. *HDI,* hydrodynamic injection; GSEA, gene set enrichment analysis; PPI, protein-protein interaction; FDR, fasle discovery rate; NES, normalized enrichment score

### PD-1 signaling sustains early proliferation and survival of HBcAg-specific CD8^+^ T cells during acute self-resolving HBV replication

To investigate the role of PD-1 expression in regulating early HBV-specific CD8^+^ T cell responses during acute HBV infection, we developed a dual adoptive transfer model using PD-1^-/-^ C93-TCRtg mice (generated by crossing PD-1 knockout mice with C93-TCRtg mice to obtain HBcAg-specific CD8^+^ T cells lacking PD-1 expression), wherein splenic CD8^+^ T cells from WT or PD-1^-/-^ C93-TCRtg mice were transferred into C57BL/6 recipients one day before HDI of the pSM2 plasmid, followed by analysis of donor CD8^+^ T cell dynamics at designated time points (**Fig. 4A**). Initial dose optimization revealed that transferring 2000 WT C93-TCRtg CD8^+^ T cells yielded significantly higher donor cell expansion in the liver compared to 500 cells at 21 dpi, while increasing to 5000 cells did not further enhance expansion (**Supplementary** Fig. 1A-C). Moreover, the adoptive transfer of 2000 cells showed no impact on HBV clearance in the model (**Fig.4B**). Therefore, we established 2000 cells as the optimal dose for subsequent experiments. Compared to WT controls, PD-1^-/-^ donor CD8+ T cells showed markedly reduced absolute numbers and frequencies in both spleen and liver at 4 and 7 dpi (**Fig. 4C**), with this early numerical deficit correlating to diminished proliferative capacity as evidenced by lower Ki-67 expression in splenic PD-1^-/-^ cells at 4 dpi (**Fig. 4D**). PD-1 deficiency had minimal impact on T_E_/T_EM_ (CD44+CD62L-) differentiation, with only a modest and transient increase in liver T_E_/T_EM_ frequencies observed at 7 dpi compared to WT controls (**Fig. 4E**). However, we observed that T-bet expression in PD-1^-/-^ cells was significantly reduced in both liver and spleen at 4 dpi and 21 dpi, while Eomes downregulation occurred in the liver at 21 dpi. PD-1 deficiency also resulted in significant downregulation of Tox expression in both liver and spleen by 21 dpi **(Fig. 4F)**, suggesting that early PD-1 signaling may sustain transcriptional programs critical for long-term T cell differentiation. Surprisingly, PD-1^-/-^ cells exhibited significantly reduced granzyme B expression by 21 dpi (**Fig. 4G**), while effector cytokine production (IFN-γ, IL-2, and TNF-α) remained largely unaffected across all timepoints, indicating that PD-1 does not suppress T cell effector function and may instead support cytotoxic killing activity (**Fig. 4H**). Taken together, these results demonstrate that PD-1 signaling sustains early proliferation and survival of HBcAg-specific CD8^+^ T cells, with its absence driving compromised cytotoxicity and decreased expression of T-bet, Eomes and Tox in HBcAg-specific CD8^+^ T cells.

**Figure 4.**
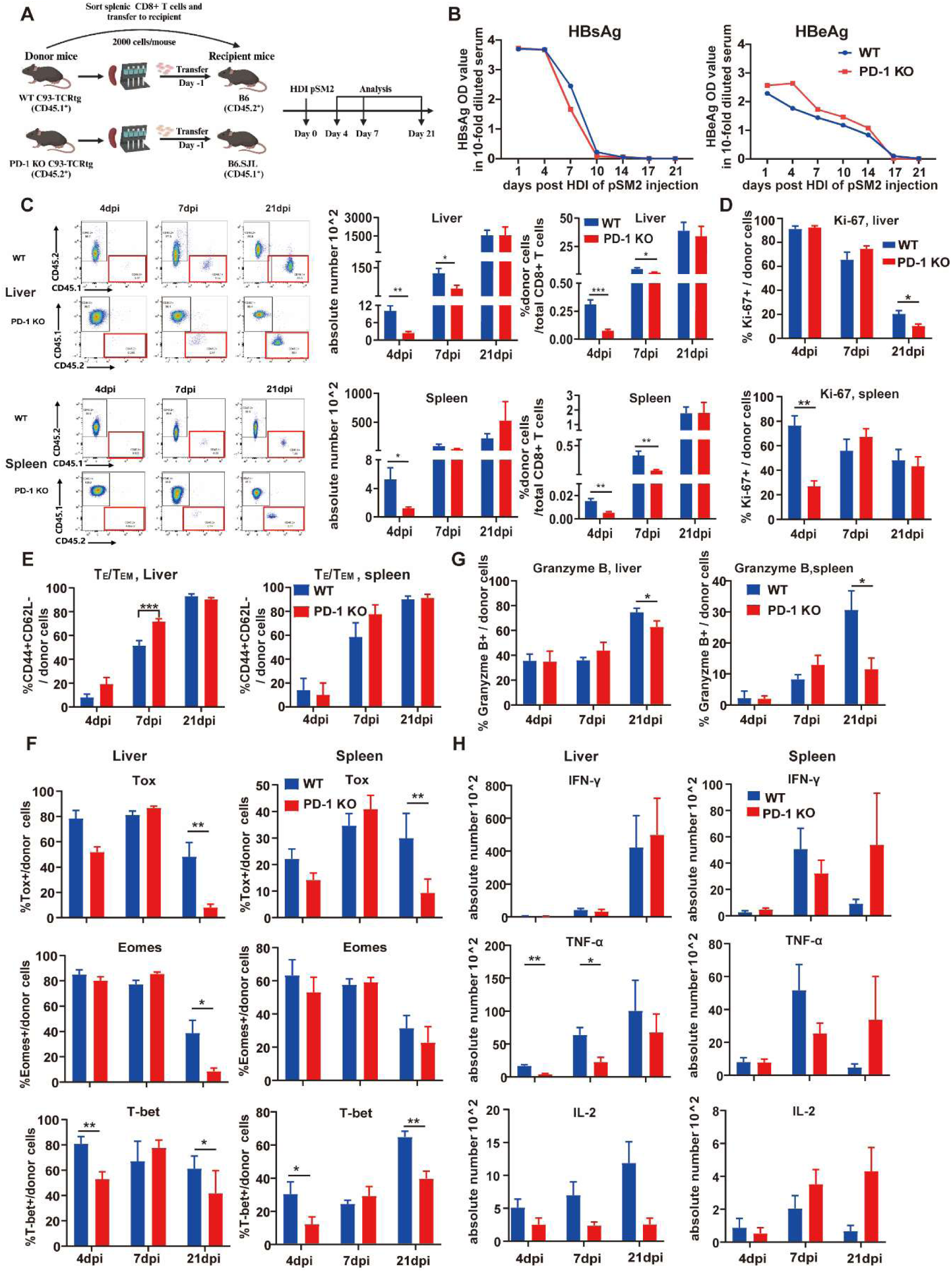
PD-1 deficiency resulted in decreased proliferation, activation, differentiation and antivirus function of HBcAg-specific CD8+ T cells upon acute-resolving HBV challenge. (A) Schematic overview of the experimental setup: 2000 purified splenic CD8+ T cells isolated from wild-type or PD-1-/-core93-TCRtg mice donor mice were transferred to C57BL/6 or C57BL/6.SJL wild-type mice at one day before HDI of pSM2 plasmid. LILs and splenocytes were collected from recipient mice at indicated time points for analysis. (B) The kinetics of serum HBsAg and HBeAg were determined. (C) The absolute number and frequency of donor CD8+ T cells in liver and spleen of recipient mice were shown. (D) The expression of proliferation marker Ki-69 on donor CD8+ T cells in liver and spleen of recipient mice were detected at indicated time points. (E) The frequency of T_E_/T_EM_ (CD44+CD62L-) on donor CD8+ T cells in liver and spleen of recipient mice were determined. (F) The expression of Eomes, T-bet and Tox on donor CD8+ T in recipient mice were displayed. (G) The frequency of degranulation marker granzyme B on donor CD8+ T cells in liver and spleen of recipient mice were detected at indicated time points. (H) LILs and splenocytes from recipient mice were stimulated with core peptide and CD28 in the presence of Brefeldin A for 3-5h. The absolute number of IFN-γ, IL-2 and TNF-α+ cells out of donor CD8+ T were displayed. Five to six mice were analyzed per group, and at least two independent experiments were performed. Data are depicted as arithmetic means ± SEM. Differences between mouse groups were analyzed using unpaired Student’s *t* tests. \**p*< 0.05, \*\**p*< 0.01, \*\*\**p*< 0.001. *HDI,* hydrodynamic injection; *LIL,* liver infiltrating lymphocytes; T_E_/T_EM,_ effector T or effector memory T.

### PD-1 deficiency exacerbates HBcAg-specific CD8^+^ T cell dysfunction in chronic HBV infection

Given the established role of PD-1 as a pivotal regulator of T cell exhaustion in chronic infections, we next investigated how PD-1 ablation impacts early development of HBV-specific CD8^+^ T cell responses in chronic HBV infection (induced by rAAV8-HBV1.3). Splenic CD8+ T cells from WT or PD-1^-/-^ C93-TCRtg mice were transferred into C57BL/6 recipients one day prior to viral challenge (**Fig. 5A**). In line with the acute infection, adoptive transfer of 2000 cells did not alter chronic viral persistence and viremia kinetics, confirming the model’s validity for T cell-intrinsic analysis (**Fig. 5B**). Unlike the robust proliferation observed in acute self-resolving infection, transferred HBcAg-specific CD8^+^ T cells exhibited minimal expansion within the first 21 days in this chronic setting, irrespective of PD-1 expression status (**Fig. 5C**). To further investigate the long-term dynamics of HBV-specific CD8^+^ T cells in chronic infection, we extended the observation period to 60 days. Intriguingly, WT donor cells exhibited delayed proliferative activity and progressive intrahepatic accumulation by 60 dpi (**Fig. 5C**), suggesting adaptive survival mechanisms under persistent antigen exposure. In stark contrast, PD-1^-/-^ donor cells completely failed to demonstrate such late-phase expansion and hepatic enrichment (**Fig. 5C**). Functional analysis further revealed that WT donor cells retained a subset capable of producing both IFN-γ and TNF-α at this chronic stage, whereas PD-1^-/-^ donor cells showed near-complete loss of IFN-γ production while maintaining limited TNF-α secretion (**Fig. 5D**). Collectively, these results demonstrating that PD-1 deficiency disrupts the delayed proliferative adaptation and effector plasticity of HBV-specific CD8^+^ T cells in chronic infection.

**Figure 5.**
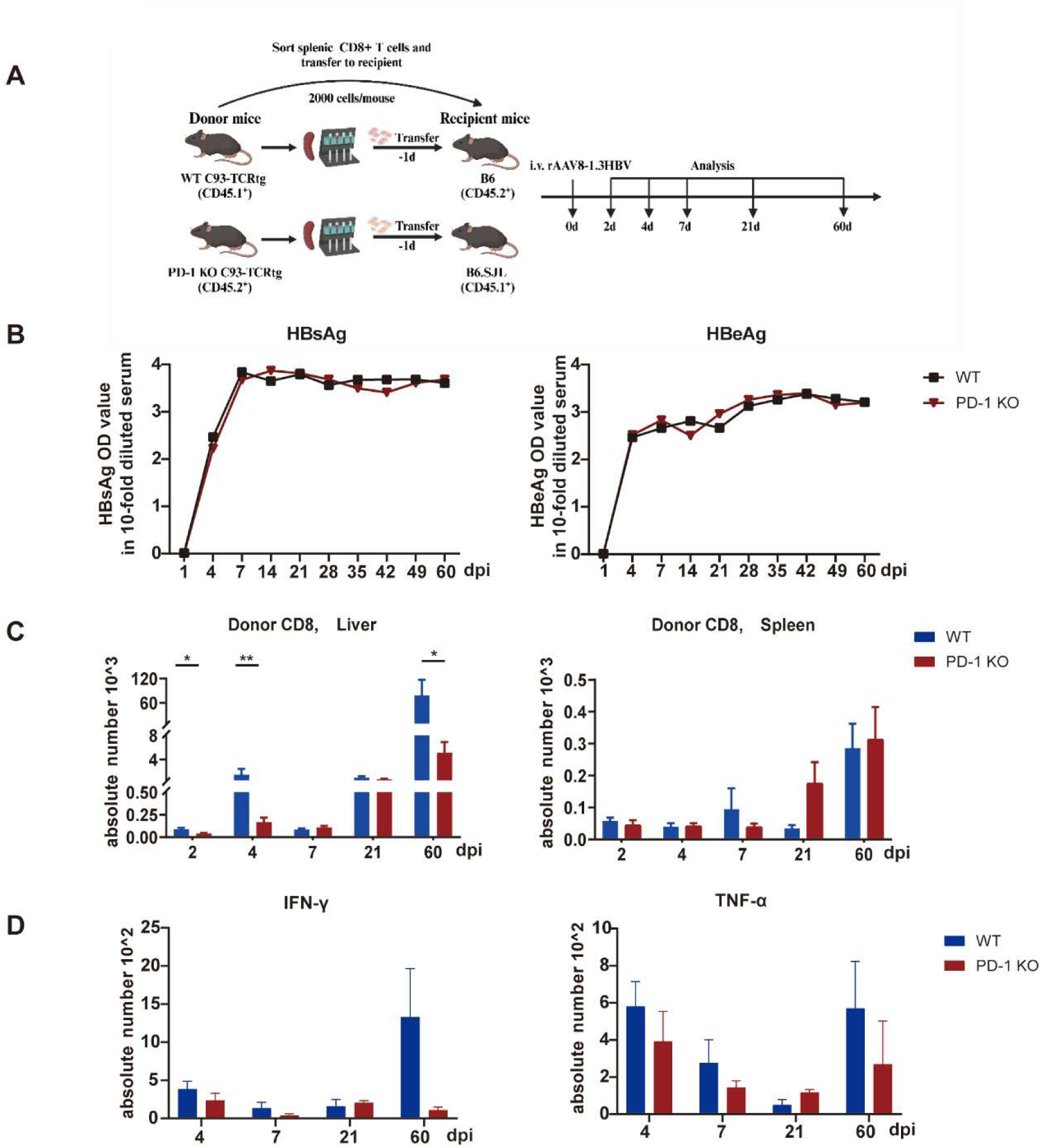
PD-1 deficiency fails to rescue the function of HBcAg-specific CD8+ T cells upon persistent HBV challenge. (A) Schematic overview of the experimental setup: 2000 purified splenic CD8+ T cells from wild-type or PD-1-/-core93-TCRtg mice donor mice were transferred to C57BL/6 or C57BL/6.SJL wild-type mice at one day before intravenous injection of rAAV8-HBV1.3. Following injection of rAAV8-1.3HBV, LILs and splenocytes as well as serum were isolated from recipient mice at indicated time points for analysis. (B)The kinetics of serum HBsAg and HBeAg were determined. (C) The absolute number and frequency of donor CD8+ T cells in liver and spleen mice were shown. (D) LILs from recipient mice were stimulated with core peptide and CD28 in the presence of Brefeldin A for 3-5h. The absolute number of IFN-γ, IL-2 and TNF-α+ cells out of donor CD8+ T were displayed. Five to six mice were analyzed per group, and at least two independent experiments were performed. Data are depicted as arithmetic means ± SEM. Differences between mouse groups were analyzed using unpaired Student’s *t* tests. \**p*< 0.05, \*\**p*< 0.01, \*\*\**p*< 0.001. *LIL,* liver infiltrating lymphocytes.

### PD-1 deficient HBcAg-specific memory CD8^+^ T cells preserve antigen-driven reexpansion and effector cytokine production

Previous studies demonstrated that CD8^+^ T cell-specific PD-1 signals are critical for establishing long-term memory during acute LCMV infection [19]. To investigate the regulatory role of PD-1 in HBV-specific CD8^+^ T cell memory responses, we adoptively transferred splenic HBcAg-specific CD8^+^ T cells (WT or PD-1 KO) into C57BL/6 recipients one day prior to hydrodynamic injection (HDI) of pSM2 plasmids. A secondary pSM2 plasmid challenge was administered at day 60 post-initial transfer (**Fig. 6A**). Serum HBsAg and HBeAg remained undetectable following the secondary challenge, confirming robust memory T cell-mediated viral control in both groups (**Fig. 6B**). Comparative analysis revealed enhanced expansion of PD-1 KO donor-derived HBcAg-specific CD8^+^ T cells at 66 dpi relative to WT counterparts (**Fig. 6C**).

**Figure 6.**
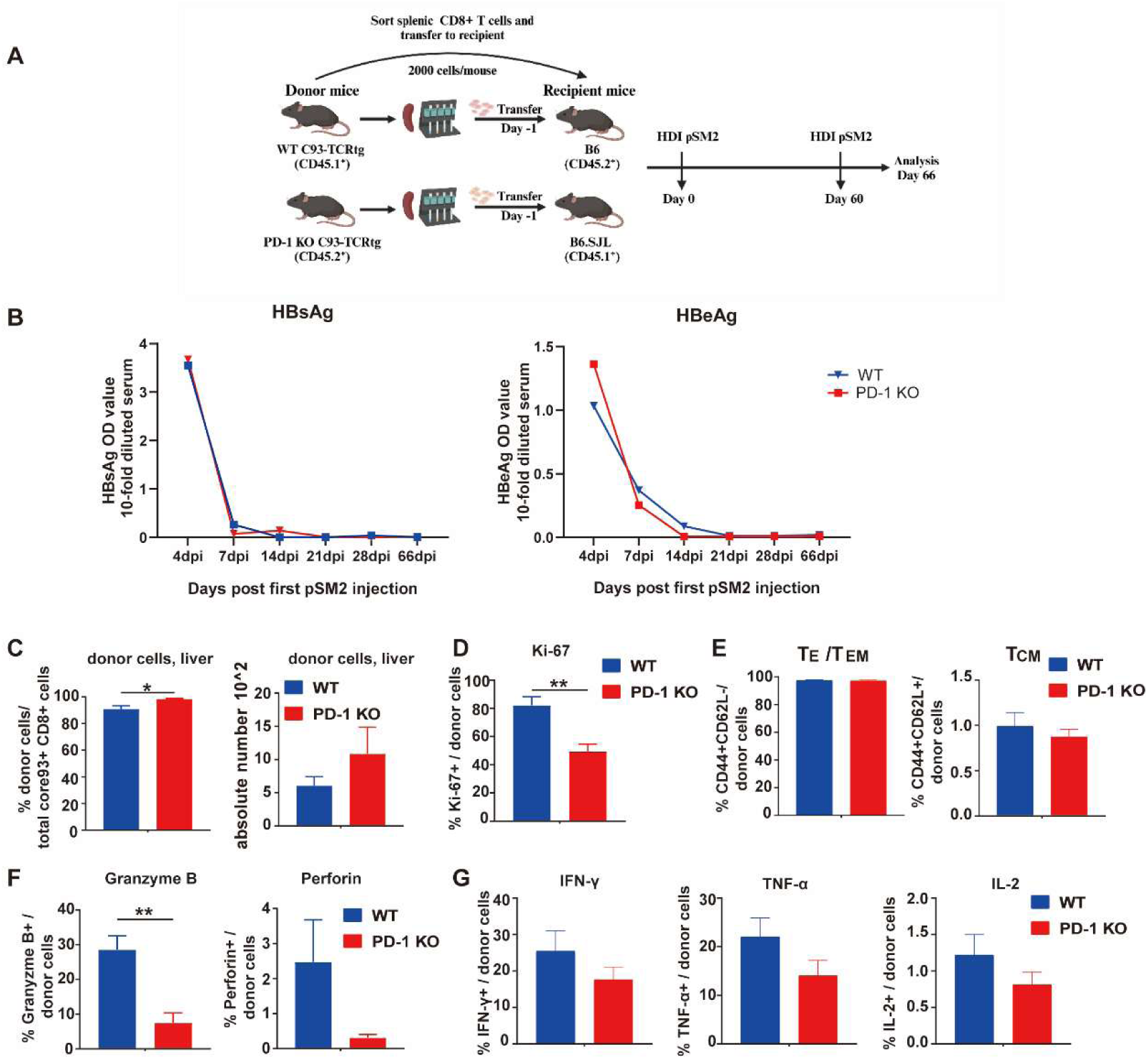
PD-1 deficiency fails to promote memory T cell response in the mouse model of secondary exposure of HBV. (A) Schematic overview of the experimental setup: 2000 purified splenic CD8+ T cells from wild-type or PD1-/-core93-TCRtg mice donor mice were transferred to C57BL/6 or C57BL/6.SJL wild-type mice at one day before HDI of pSM2 plasmid. At 60 dpi, mice were injected with pSM2 plasmid by HDI for second time. LILs and splenocytes were isolated from recipient mice at day 6 post second injection of pSM2 plasmid. (B) The kinetics of serum HBsAg and HBeAg were determined. (C) The frequency and absolute number of donor CD8+ T cells in liver recipient mice were shown at 66 dpi. (D) The frequency of Ki-67 of donor CD8+ T cells between two groups was shown. (E) The differentiation of donor CD8+ T cells on recipient mice were determined. (F) The expression of granzyme B and perforin on donor cells between two groups was shown. (G) LILs and splenocytes from recipient mice were stimulated with CD3 and CD28 in the presence of Brefeldin A for 3-5h. The frequency of IFN-γ, IL-2 and TNF-α+ cells out of donor CD8+ T were displayed. Five to six mice were analyzed per group. Data are depicted as arithmetic means ± SEM. Differences between mouse groups were analyzed using unpaired Student’s *t* tests. \**p*< 0.05, \*\**p*< 0.01. *dpi,* days post-injection; *HDI,* hydrodynamic injection; *KO*, knockout.

Intriguingly, despite this increased frequency and absolute number of PD-1 KO cells, they exhibited reduced expression of the proliferation marker Ki-67 at 66 dpi (**Fig. 6D**). Critically, however, analysis of effector T cell differentiation status revealed no significant differences between PD-1 KO and WT donor cells in the frequencies of T_E_/T_EM_ and central memory (T_CM_) subsets (**Fig. 6E**). Furthermore, functional profiling of HBV-specific CD8^+^ T cells demonstrated that PD-1 KO cells exhibited reduced granzyme B and perforin expression, but produced comparable levels of IFN-γ, TNF-α, and IL-2 compared to WT cells (**Fig. 6F and G**). These collective findings suggest that PD-1 ablation does not substantially compromise the memory responsiveness or cytokine secretory capacity of HBV-specific CD8^+^ T cells, despite transient alterations in proliferative kinetics and cytolytic molecule expression.

### PD-1 deficiency promotes activation-induced apoptosis in HBV-Specific CD8^+^ T cells

Given that PD-1 ablation consistently reduced HBV-specific CD8^+^ T cell numbers and proliferative capacity in both acute and chronic infection models, we sought to delineate the underlying mechanisms. We established an in vitro stimulation system mimicking antigen-specific activation, where splenocytes from WT or PD-1^-/-^ C93-TCRtg mice were CFSE-labeled and stimulated with HBc peptide plus anti-CD28 (**Fig.7A**). Consistent with our in vivo observations, PD-1^-/-^ HBcAg-specific CD8^+^ T cells exhibited markedly impaired proliferation and reduced IFN-γ/TNF-α production compared to WT counterparts over 72–96 hours (**Fig. 7B-C**), confirming the cell-intrinsic role of PD-1 in sustaining T cell expansion and effector function. Next, we performed transcriptomic profiling on cells harvested pre-stimulation and at multiple post-stimulation timepoints (24–96 h) via RNA sequencing (**Fig. 8A**). At 48h post-stimulation, PD-1 KO CD8^+^T cells displayed the largest differentially expressed genes among the four time points compared to the WT CD8^+^ T cells, identifying 3778 upregulated and 3916 downregulated genes (**Fig.8B**). At 0h, there were 11 differentially expressed genes associated with cell cycle, replication and death (**Fig.8C**). Notably, the cell cycle pathway was consistently enriched among the top 15 differentially expressed KEGG pathways across all four time point between WT and PD-1 KO CD8^+^ T cells (**Fig.8D**). Consistently, PD-1 KO HBcAg-specific CD8^+^ T cells exhibited trend of enrichment of apoptosis gene signature at day 0, 24h and 72h after stimulation (**Fig.8E**). Mechanistic dissection demonstrated that PD-1 deficiency significantly upregulated pro-apoptotic effectors (*Bax, NOXA, Apaf1, Casp3*) while suppressing the anti-apoptotic regulator *Bcl2*, with maximal effects at 48h post-stimulation (**Fig. 8F**). This gene signature indicates that PD-1 loss might exacerbate activation-induced apoptosis, mechanistically explaining the progressive numerical decline of HBV-specific CD8^+^ T cells in vivo. Collectively, PD-1 restrains activation-induced apoptosis through transcriptional control of apoptotic machinery, thereby preserving the survival and functional persistence of antiviral CD8^+^ T cells.

**Figure 7.**
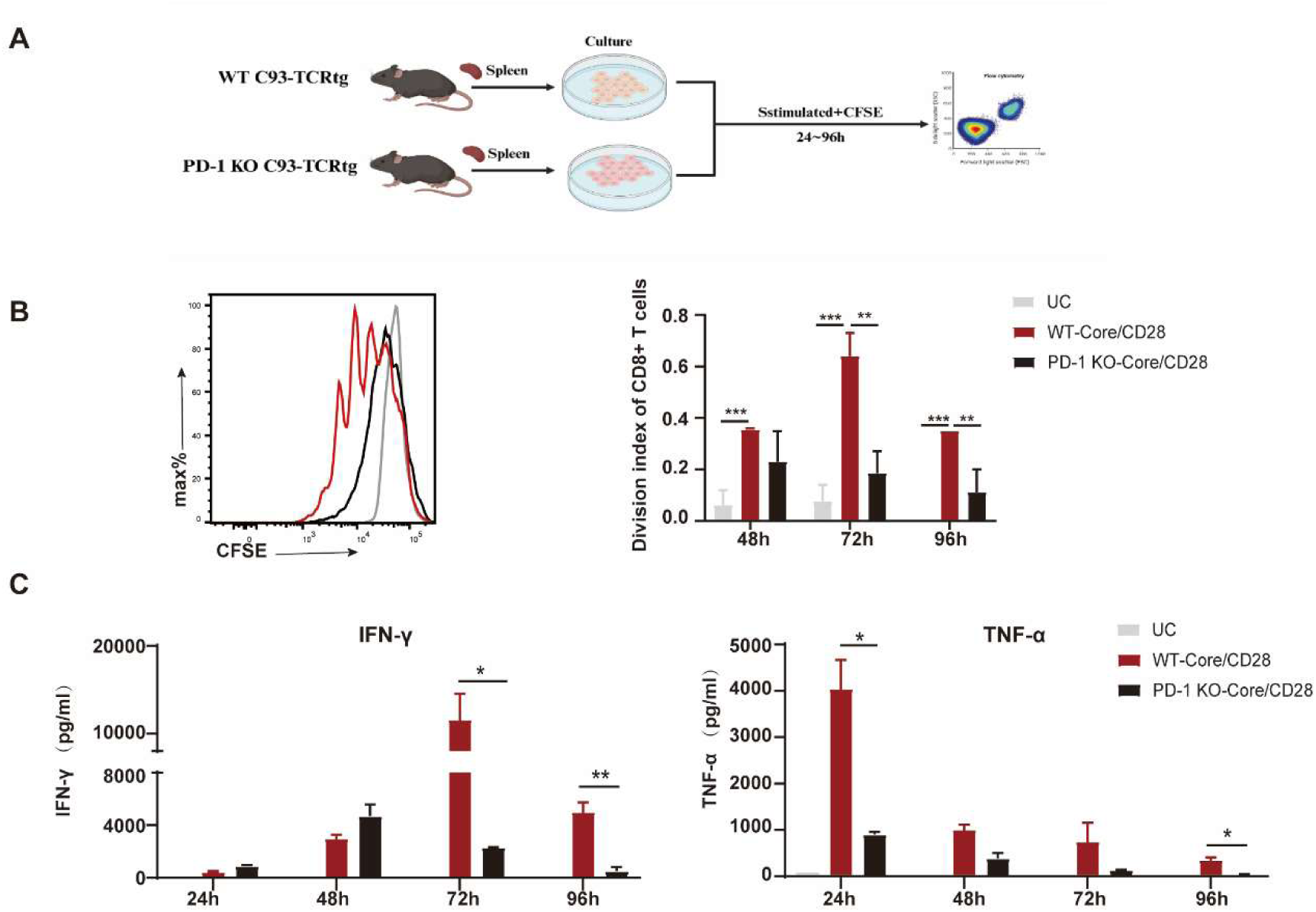
PD-1 deficiency results in decreased proliferation, effector function of HBcAg-specific CD8+ T cells *in vitro*. (A) Schematic overview of the experimental setup: splenocytes isolated from wild-type or PD-1^-/-^ core93-TCRtg mice were labled with CFSE and subsequently stimulated with core peptide and CD28 for 24-96 h. The proliferation of CD8+ T cell were detected by flow cytometry. Secretion of IFN-γ and TNF-α in the supernatant were detected at indicated time points. (B) The proliferation and the division index of CD8+ T cells were shown. (C) The expression of *IFN-γ* and TNF-α in the supernatant were determined. Data are depicted as arithmetic means ± SEM. The statistical differences among multiple groups were analyzed by one-way ANOVA followed by Turkey’s multiple comparisons test. \**p*< 0.05, \*\**p*< 0.01, \*\*\**p*< 0.001. *CFSE*, Carboxyfluorescein succinimidyl ester; *IFN-γ*, interferon-gamma; *TNF-α*, tumor necrosis factor-alpha.

**Figure 8.**
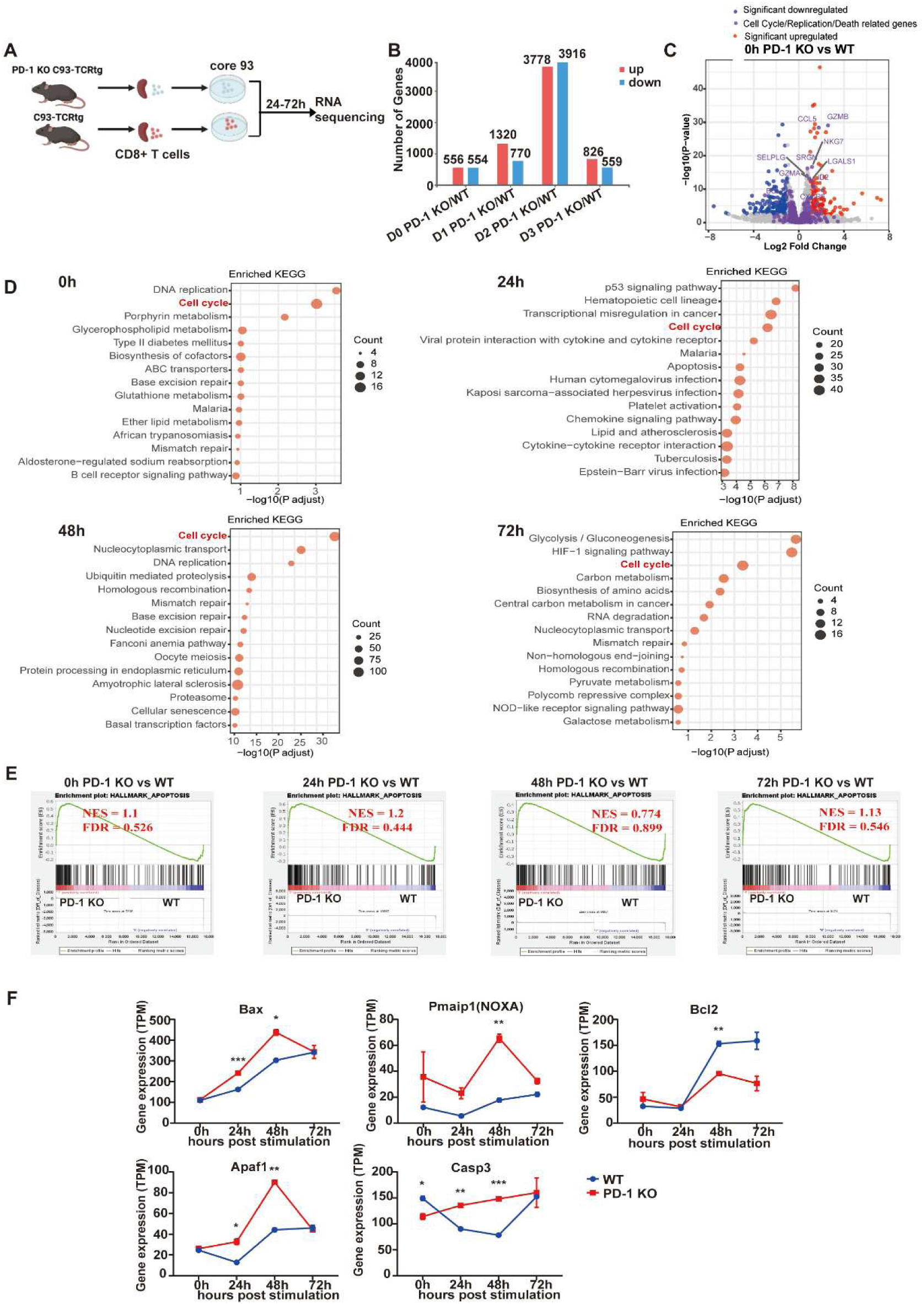
PD-1 deficiency results in a significant increase in the activation of apoptosis signaling pathway in T cells. (A) Schematic overview of the experimental setup: CD8+T cells isolated from the spleen of wild-type or PD-1^-/-^ core93-TCRtg mice were stimulated with core peptide and CD28 for 24-72h. Cells were collected for RNA sequencing detection before and after stimulation at different time points. (B) The number of differentially expressed genes between two groups at different time points was shown. (C) Volcano plots showed differentially expressed genes between two groups at 0h. The violets dots showed cell cycle, replication and death related genes. (D) Top 15 KEGG enriched terms for significant regulated genes between two groups at different time points were shown. (E) GSEA analysis showed enrichment of an apoptosis signature (Hallmark apoptosis) in CD8+ T cells from PD-1 KO versus WT mice at different time points. (F) Dynamic of apoptosis signaling markers were shown. Two samples per group were shown. Two samples were used in each group at each time point. Data are depicted as arithmetic means ± SEM. Differences between mouse groups were analyzed using unpaired Student’s *t* tests. \**p*< 0.05, \*\**p*< 0.01, \*\*\**p*< 0.001. KO, knock out, FDR, fasle discovery rate; NES, normalized enrichment score

## Discussion

This study delineates a critical and context-dependent role for PD-1 in shaping HBV-specific CD8^+^ T cell responses during acute infection. Integrating our key findings, we demonstrate that: (i) PD-1 exhibits sustained and pronounced upregulation on effector-phenotype HBV-specific CD8^+^ T cells in acute but not chronic HBV replication; (ii) PD-1^+^ CD8^+^ T cells display enhanced proliferative activity (Ki-67), cytotoxic potential (granzyme B), and cytokine production compared to PD-1^-^ counterparts in AR infection; (iii) Genetic ablation of PD-1 significantly impairs the early expansion and survival of HBV-specific CD8+ T cells in both acute and chronic settings, linked to dysregulated cell cycle progression and enhanced apoptosis; and (iv) Despite impaired expansion, PD-1 deficiency does not suppress effector cytokine production (IFN-γ, TNF-α, IL-2) in acute infection but exacerbates functional exhaustion in chronicity.

Our observation of robust PD-1 upregulation on functional effector CD8^+^ T cells during acute self-resolving HBV infection aligns with findings in acute Friend virus and HCV infections, where PD-1 marks activated cytotoxic effectors rather than exhaustion [15, 20]. The stark contrast in PD-1 dynamics between AR and CH models underscores its divergent roles contingent on infection outcome. In AR, sustained PD-1 expression likely reflects persistent antigen presentation by dendritic cells (DCs) within lymphoid and hepatic microenvironments, driving ongoing T cell activation [12, 13]. The predominance of a T_E_/T_EM_ phenotype (CD44+CD62L-) within the PD-1+ compartment, coupled with heightened granzyme B and cytokine production, positions PD-1 as a co-regulator of effector differentiation in acute HBV resolution. This contrasts with chronic settings where persistent antigen and inflammation drive PD-1’s canonical inhibitory role [21, 22].

The most striking finding was the requirement of PD-1 for optimal early expansion of HBV-specific CD8^+^ T cells. PD-1 KO cells exhibited profound deficits in proliferative capacity (reduced Ki-67 and CFSE dilution) and numerical accumulation in spleen and liver during acute infection. This defect was mechanistically linked to transcriptional dysregulation of cell cycle and apoptosis pathways. PD-1 deficiency triggered upregulation of pro-apoptotic genes (Bax, Noxa, Casp3) and downregulation of anti-apoptotic Bcl2, particularly upon antigen encounter. This aligns with studies showing PD-1 engagement can promote survival in activated T cells by modulating PI3K/Akt signaling and glucose metabolism[19]. Our data thus reveal a T cell-intrinsic pro-survival function for PD-1 during the initial clonal burst, potentially counteracting activation-induced cell death (AICD) to ensure sufficient effector pool size for viral control. This challenges the simplistic view of PD-1 solely as an inhibitor of early T cell responses.

Importantly, while PD-1 ablation crippled expansion and granzyme B expression, it did not suppress the ability of HBV-specific CD8^+^ T cells to produce IFN-γ, TNF-α, or IL-2 during acute infection. This dissociation between expansion/cytotoxicity and cytokine production suggests PD-1 regulates distinct functional modules. The significant downregulation of T-bet (early and late), Eomes (late), and Tox (late) in PD-1 KO cells points to a role for PD-1 in sustaining transcriptional networks crucial for effector differentiation and potentially memory formation, consistent with its role in regulating mTOR-dependent metabolic pathways in memory precursors [19].

In chronic HBV persistence, the absence of PD-1 failed to prevent clonal depletion and instead exacerbated dysfunction. PD-1 KO cells lacked the delayed proliferative adaptation seen in WT cells and exhibited a profoundly exhausted phenotype (loss of IFN-γ production). This mirrors findings in chronic LCMV, where PD-1 deficiency accelerates terminal exhaustion [23], and highlights that the functional consequence of PD-1 loss is dictated by the inflammatory and antigenic milieu. The inability of PD-1 KO cells to persist long-term in chronic infection, despite initial activation, underscores its non-redundant role in maintaining T cell fitness under persistent antigen pressure.

Our findings have translational implications. Firstly, they suggest that therapeutic PD-1/PD-L1 blockade initiated very early in acute HBV infection might inadvertently impair the crucial initial expansion of virus-specific CD8^+^ T cells, potentially compromising viral clearance. Secondly, the observation that HBV-specific CD8^+^ T cell exhaustion and clonal depletion occur, even in the absence of PD-1, in chronic infection (**Fig 5C-D**) indicates that alternative inhibitory pathways (e.g., CTLA-4, TIGIT, LAG3, identified in our transcriptome data) or intrinsic factors can drive exhaustion. This necessitates combinatorial targeting strategies for restoring immunity in CHB, as suggested by our prior woodchuck studies [8]. The potential for PD-1 blockade to cause bystander memory T cell attrition, as reported in chronic LCMV[19], warrants investigation in HBV settings.

Our data demonstrate that genetic ablation of PD-1 does not significantly impair the recall response or cytokine production of HBV-specific memory CD8^+^ T cells upon secondary challenge (Fig. 6). This finding contrasts with prior reports in acute LCMV and influenza infections, where constitutive PD-1 deficiency led to defective memory CD8+ T cell formation and stability[19, 24]. We propose several non-exclusive explanations for this dichotomy: (i) HBV exhibits distinct tropism (hepatotropic) and replication dynamics compared to systemic (LCMV) or respiratory (influenza) viruses. The liver’s tolerogenic microenvironment, characterized by high PD-L1 expression on sinusoidal endothelial cells and immunosuppressive cytokines (e.g., IL-10, TGF-β), may reshape PD-1 function. In HBV resolution, sustained PD-1+ effector T cells maintain functionality (Fig. 2), suggesting compensatory pathways may preserve memory differentiation independent of PD-1. (ii) Influenza and LCMV models show prolonged antigen presentation (30+ days), requiring PD-1 to temper excessive early proliferation for memory precursor survival. In contrast, acute-resolving HBV in our model features rapid clearance (HBsAg-negative by 21 dpi), potentially reducing dependence on PD-1 for memory cell survival. This aligns with transient PD-1 blockade studies in influenza where early inhibition did not disrupt memory [24]. (iii) Transcriptomic data reveal upregulation of alternative exhaustion markers (CTLA-4, TIGIT, LAG-3) on PD-1^+^ HBV-specific CD8^+^ T cells (Fig. 3E). In chronic settings, PD-1 deficiency failed to prevent exhaustion (Fig. 5), implying redundancy among inhibitory receptors in HBV infection. Such pathways may sustain memory homeostasis in the absence of PD-1. The context-dependent role of PD-1—pro-survival in acute HBV effectors yet dispensable for memory—highlights its functional plasticity across infections. This underscores the need to evaluate checkpoint regulators in disease-specific milieus, particularly for therapies targeting PD-1 in HBV-related disorders.

Certain limitations of our study should be considered. Firstly, our HBV hydrodynamic injection model, while valuable, differs from natural infection routes; validating key findings in patient-derived samples or more physiological models is crucial. Secondly, the adoptive transfer system, though controlled, bypasses natural T cell priming and homing dynamics.

In summary, this study redefines PD-1’s role in acute HBV infection. Rather than merely constraining effector function, PD-1 is essential for the early expansion and survival of HBV-specific CD8^+^ T cells, acting as a safeguard against activation-induced apoptosis. Its absence leads to crippled clonal expansion and impaired cytotoxicity without suppressing cytokine potential acutely, while promoting exhaustion chronically. These context-dependent functions position PD-1 as a critical rheostat calibrating the magnitude and quality of the antiviral CD8^+^ T cell response, with significant implications for immunotherapeutic strategies targeting this pathway in hepatitis B.

## Acknowledgements

We thank Dr.Yaojie Liang from Zhejiang Provincial Cancer Hospital for the technical help of GSEA analysis.

## Authors’ contributions

JL and YD conceived and designed the experiments. XZ, WP, ZL, HH and XY performed the experiments. XZ, WP, ZL and YD analyzed the data. KS and ML, UD, GZ, XZ, DY and PK provide technical guidance for the study. YD and JL wrote the manuscript. KS, ML, UD, GZ, XZ, DY, PK, and JL revised the manuscript. All authors read and approved the final version.

## Data Availability statement

All data generated or analyzed during this study are included in this article and its additional files. The data used in the current study are available from the corresponding author on reasonable request.

## Abbreviations

AR: acute self-resolving HBV replication
AICD: activation-induced cell death
CHB: chronic hepatitis B
CH: chronic HBV replication
CTLA4: cytotoxic T-lymphocyte associated protein 4
C93-TCR-tg: Core93 TCR-transgene
DCs: dendritic cells
ELISA: enzyme-linked immunosorbent assay
FDR: false discovery rate
FV: Friend virus
GSEA: Gene set enrichment analysis
HBV: Hepatitis B virus
HBsAg: hepatitis B surface antigen
HBeAg: hepatitis B e antigen
HCV: hepatitis C virus
HDI: hydrodynamic injection
IFN-γ: interferon-gamma
IL-2: interleukin-2
KO: knockout
LCMV: lymphocytic choriomeningitis virus
LIL: liver-infiltrating lymphocyte
NS: normal saline
PD-1: programmed cell death protein 1
PD-L1: programmed cell death protein ligand1
PBMCs: peripheral blood mononuclear cells
PBS: phosphate-buffered saline
WHV: woodchuck hepatitis virus
WHsAg: woodchuck hepatitis surface antigen
SEM: mean ± standard error
SPF: specific pathogen-free
TCR: T cell receptor
WT: Wild type
T_E_/T_EM,_: Effector and effector memory T cell
TNF-α: tumor necrosis factor-alpha.

