## supplementary for "PD-1 signaling is essential for the early accumulation of HBV-specific CD8+ T cells during HBV infection"

**PD-1 is required for the early expansion of HBV-specific CD8+ T cells and does not suppress their effector function during the acute phase of HBV infection**

**Xiaoqing Zeng*[1, 2], Wen Pan*[1, 2], Ziwei Li[1, 2] , Zhaoli Liu[3], Hongming Huang[1, 2], Xuecheng Yang[1, 2], Kathrin Sutter[4,5], Mengji Lu[4], Ulf Dittmer[4], Gennadiy Zelinskyy[4], Xin Zheng[1, 2], Dongliang Yang[1, 2], Patrick T.F. Kennedy[6], Yanqin Du[#1, 2], Jia Liu[#1, 2]**

1. Department of Infectious Diseases, Union Hospital, Tongji Medical College, Huazhong University of Science and Technology, Wuhan 430022, China
2. Institute of Infectious Diseases and Immunity, Union Hospital, Tongji Medical College, Huazhong University of Science and Technology, Wuhan, 430022, China
3. Center China Subcenter of National Center for Cardiovascular Diseases, Zhengzhou 451460, China
4. Institute for Virology, University Hospital Essen, University of Duisburg-Essen, Essen 45147, Germany
5. Institute for the Research of HIV and AIDS-associated diseases, University Hospital Essen, University of Duisburg-Essen, Essen 45147, Germany
6. Barts Liver Centre, Blizard Institute, Barts and the London School of Medicine and Dentistry, Queen Mary University of London, London, UK.

**Table of contents**

**Supplementary Table 1**

**Supplementary figure 1**

**Supplementary Table 1. Antibodies used for flow cytometry**

| Antibody | Color | Source |
|---|---|---|
| Anti-mouse CD45.1 | PE | BioLegend |
| Anti-mouse CD45.1 | FITC | BioLegend |
| Anti-mouse CD45.2 | PE | BioLegend |
| Anti-mouse CD45.2 | FITC | BioLegend |
| Anti-biotin | PE | Miltenyi Biotec |
| Anti-mouse CD4 | APC-cy7 | BioLegend |
| Anti-mouse PD-1 | BV421 | BD Bioscience |
| Anti-mouse CD8 | FITC | BD Bioscience |
| Anti-mouse CD43 | PE-cy7 | BD Bioscience |
| Anti-mouse CD44 | APC | BD Bioscience |
| Anti-mouse CD62L | PE-cy7 | BD Bioscience |
| Anti-mouse CD69 | Percp-cy5.5 | BD Bioscience |
| Anti-mouse IFN-γ | APC | BD Bioscience |
| Anti-mouse IL-2 | percp cy5.5 | BioLegend |
| Anti-mouse TNF-α | FITC | BD Bioscience |
| Anti-mouse Granzyme B | PE | BioLegend |
| Anti-mouse Ki-67 | PB | BD Bioscience |
| Anti-mouse TOX | PE | BD Bioscience |
| Anti-mouse T-bet | APC | BD Bioscience |
| Anti-mouse Eomes | PE-cy7 | BD Bioscience |

Supplementary figure 1

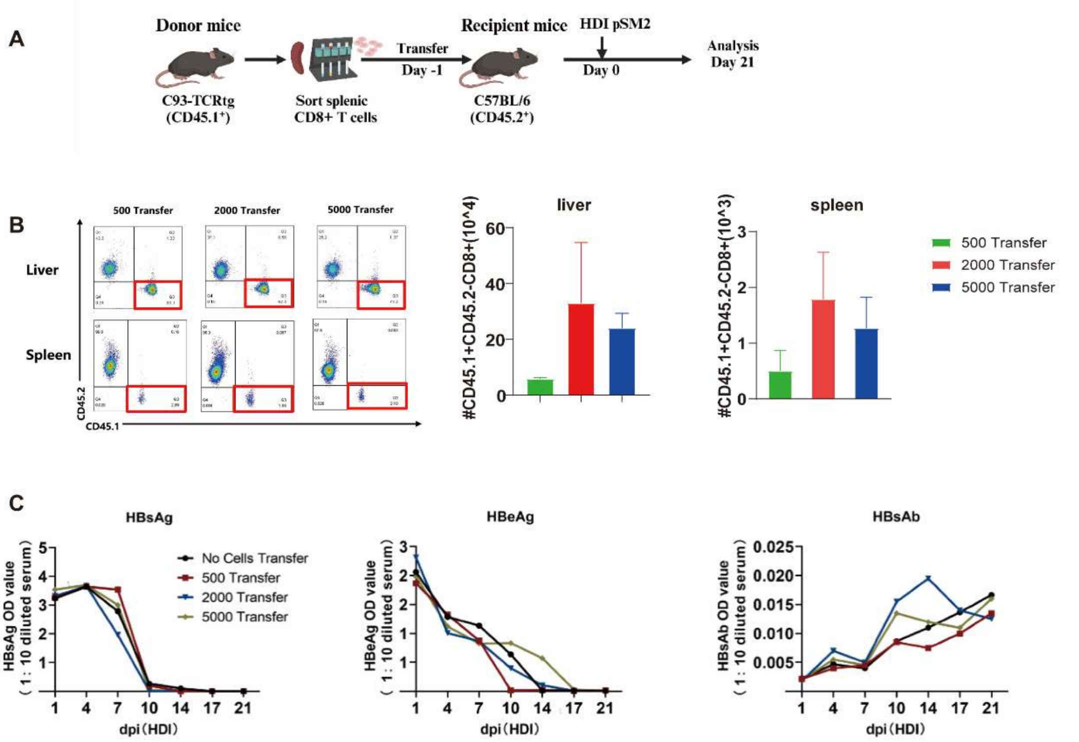


**Supplementary figure 1. The effects of different number of transferred HBcAg-specific CD8+ T cell on HBV replication.** (A) Schematic overview of the experimental setup: different number of purified splenic CD8+ T cells from CD45.1+core93-TCRtg mice were transferred to CD45.2+ C57BL/6 wild-type mice at one day before HDI of pSM2 plasmid. Then mice were HDI of 10 μg pSM2 plasmid at day 0 and mice were sacrificed at 21 dpi. (B) The kinetics of serum HBsAg, HBeAg and HBcAb were determined. (C) The absolute number of CD45.1+ HBcAg-specific T cells on recipient mice were detected at day 21 dpi. Data are depicted as arithmetic means ±SEM. The statistical differences among multiple groups were analyzed by one-way ANOVA followed by Turkey's multiple comparisons test. The statistical differences of viral indicators were analyzed by repeated measures analysis. Five to six mice were analyzed per group. *dpi,* days post-injection; *HDI,* hydrodynamic injection.
